# The hair cycle underlies regulation of Langerhans cell renewal

**DOI:** 10.1101/832774

**Authors:** Benjamin Voisin, Dimitri Chartoire, Caroline Devouassoux, Christine Kowalczyk-Quintas, François Clauss, Frédéric Lezot, Pascal Schneider, Vincent Flacher, Christopher G. Mueller

**Affiliations:** CNRS UPR 3572 / I2CT, Laboratory of Immunology, Immunopathology and Therapeutic Chemistry / Laboratory of Excellence MEDALIS, Institut de Biologie Moléculaire et Cellulaire, Strasbourg, France; Department of Biochemistry, University of Lausanne, Epalinges, Switzerland; INSERM UMR1109 Osteoarticular and Dental Regenerative Nanomedicine, Faculté de Chirurgie Dentaire, Pôle de Médecine et de Chirurgie Bucco-Dentaires, Strasbourg, France; UFR Médecine et Techniques Médicales, Pôle Os - Articulations - Chirurgie Plastique, Nantes, France

## Abstract

In the epidermis, Langerhans cells (LCs) provide an essential link between the innate and adaptive immune systems. They self-renew in situ and continuously transport antigen from skin to lymph node (LN) T cells in the steady state. The cyclic renewal of hair follicles (HF) causes profound alterations in the cutaneous microenvironment, however little is known about its impact on LC homeostasis. Here we show that mouse LCs developed normally in the absence of hair but perceived critical transition periods in the hair cycle. LCs underwent a proliferation burst during the HF growth phase (anagen). Reinitiation or abolishment of anagen as well as loss of the HF had direct consequences on LC self-renewal. Because dividing LCs were found close to the anagen HF, we searched for the proliferative signal within this structure and identified increased *Il34* expression by HF stem cells and their progeny. Inhibition of the IL-34 receptor CSF-1R at the onset of anagen completely and specifically blocked LC proliferation. Altogether, our findings demonstrate that the hair cycle directly oversees LC self-renewal and migration.

## INTRODUCTION

Organized in a dense network within the epidermis, Langerhans cells (LCs) are the outermost sentinels of the skin immune system (**Kaplan, 2017;Doebel *et al.,* 2017**). Their location allows them to efficiently collect antigens from keratinocytes, commensal or pathogenic microorganisms, or topically-applied chemicals. When activated by environmental danger signals, LCs migrate into skin-draining LNs where they present antigens to T cells. In the steady state, the traffic occurs continuously but at a lower frequency. LCs that reach LNs without prior exposure to danger signals are thought to contribute to immune tolerance (**Steinman and Nussenzweig, 2002;Flacher *et al.,* 2014;Idoyaga *et al.,* 2013;Seneschal *et al.,* 2012**). Recently, molecular mechanisms governing this spontaneous emigration have been revealed (**Bobr *et al.,* 2012;Zahner *et al.,* 2011**), although further investigations on their initiation are still needed. Development of the LC network requires precursors derived from the yolk sac or the fetal liver and recruited into the embryonic skin (**Hoeffel *et al.,* 2012;Schulz *et al.,* 2012**). Shortly after birth, these precursors differentiate into ***bona fide*** LCs and undergo intense but transient proliferation to take up residence in the newly formed epithelium (**Chorro *et al.,* 2009**). In adult mice, the integrity of the network is maintained by a low-rate self-renewal (**Giacometti and Montagna, 1967;Czernielewski *et al.,* 1985;Merad *et al.,* 2002**), likely originating from a specialized LC subset endowed with a higher proliferative capacity (**Ghigo *et al.,* 2013**). This unique homeostatic maintenance of LCs together with their constant traffic to the draining LNs raises important questions regarding the existence of local and/or temporal control mechanisms. The hair follicle (HF) is a complex multilayered formation that extends from the epidermis deep into the dermis, and integrates sebaceous glands (**Schneider *et al.,* 2009;Hsu *et al.,* 2014**). It protects mammals against extreme temperatures, UV light or physical trauma. However, at the same time, the HF provides a niche for microorganisms that can challenge the immune system (**Polak-Witka *et al.,* 2019**). HF morphogenesis draws its origins from the interaction between the embryonic ectoderm and the underlying mesoderm and is completed two weeks after birth (**Schneider *et al.,* 2009**). A key feature of HF is its cyclic renewal, which allows for the replacement of damaged hair shaft and seasonal adjustments to the fur coat. The cycle is subdivided in three main stages: growth (anagen), regression (catagen) and resting (telogen) (**Schneider *et al.,* 2009**). These phases are under the control of complex regulatory mechanism entailing periodic activation and quiescence of HF-associated stem cells (**Hsu *et al.,* 2014**). The first cycle is synchronized for most HFs until the second telogen. Later on, HFs are uncoupled and their renewal occurs with variable kinetics within different areas of the skin (**Hodgson *et al.,* 2014;Plikus *et al.,* 2011**). It has been recognized that the hair cycle impinges on skin physiology a number of important changes (**Stenn and Paus, 2001**). By using HF synchronization mouse models, studies have demonstrated variations in the number or activation of perifollicular macrophages, dendritic epidermal T cells (DETCs) and mast cells (**Castellana *et al.,* 2014;Paus *et al.,* 1998;Westgate *et al.,* 1991;Hashizume *et al.,* 1994;Kumamoto *et al.,* 2003**). It has been known for a long time that LCs associate with the HF (**Breathnach, 1963;Moresi and Horn, 1997;Christoph *et al.,* 2000**), particularly in the non-cycling distal portion and the nearby sebaceous glands (**Haid *et al.,* 2015**) that is most exposed to trauma and infection. When acute, inflammation-induced LC emigration requires the recruitment and differentiation of precursors to replenish the network (**Katz *et al.,* 1979;Ginhoux *et al.,* 2006**), HFs have been depicted as a portal to blood-derived precursors (**Nagao *et al.,* 2012**) and as a niche for keratinocytes that support TGF-⍰-driven differentiation (**Mohammed *et al.,* 2016**). Finally, deciphering the immunosurveillance of HFs is particularly important because they have been proposed as a privileged route of entry of bioactive molecules (**Knorr *et al.,* 2009**).

In spite of these elements, the effect of the periodic activation of hair renewal and regression on LC biology remains unknown. By establishing temporal associations with the synchronized hair cycle or by its physical and genetic manipulation, we present evidence that the hair cycle regulates LC self-renewal by CSF-1R engagement. These findings show that the dynamic changes in skin physiology elicited by the hair cycle can have a major impact on the cutaneous immune system.

## RESULTS

### The Langerhans cell network develops in the absence of hair follicles

The development of the LC network coincides with HF formation between embryonic day 14.5 and post-natal day 15 (**Schulz *et al.,* 2012;Schmidt-Ullrich and Paus, 2005**), and immunofluorescence staining of hairy (tail skin) epidermal sheets revealed a close association between LCs and the HF infundibulum (**Fig. 1A**). In light of this spatiotemporal relationship between LCs and HFs, we first asked whether formation of the LC network is dependent on HF morphogenesis. Among the different mouse models lacking hair, mice deficient for epithelial morphogen ectodysplasin-A (EDA), a TNF-family member, or its receptor are particularly relevant because they display an embryonic deficiency in HFs (**Gruneberg, 1971;Headon and Overbeek, 1999**). The *Tabby* mouse is a natural mutant of EDA and is completely devoid of HFs on the tail and behind the ears because of a complete lack of morphogenesis (**Mikkola, 2008**). Immunofluorescence staining for MHC-II in hairless tail epidermis of *Tabby* mice revealed the presence of a LC network comparable to that of wild-type mice. (**Fig. 1B**). Therefore, HF morphogenesis is dispensable for LC development and residence in the epithelium.

**Figure 1:**
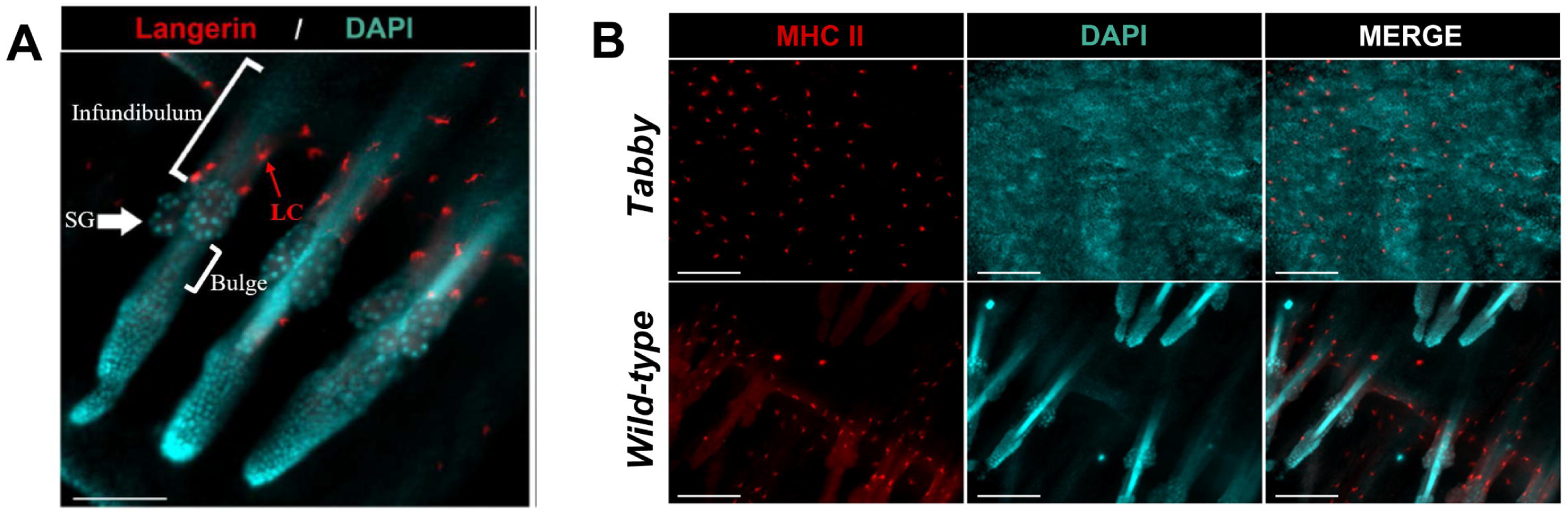
Langerhans cell network formation is independent of hair follicle morphogenesis. **(A)** Epidermal sheets comprising complete HFs from C57BL/6 tail skin were stained for Langerin (red) and DAPI (blue). The image shows LCs in an epidermal network that comprises the upper HF (infundibulum, outlined by a bracket). The arrow points to a sebaceous gland. Scale bar: 100μm. **(B)** Epidermal sheets obtained from tail skin of hairless *(Tabby)* or wild-type littermate control mice were stained for MHC class II (red) and DAPI (blue). Scale bar: 100μm.

### A Langerhans cell proliferation burst occurs concomitantly with the anagen hair cycle phase

To assess the impact of the hair cycle on LC renewal we took advantage of the synchronized first postnatal hair cycle of mouse back skin (**Fig. 2A**) (**Schneider *et al.,* 2009**). At critical time points of the cycle, back skin was processed with dispase II to allow a clean separation of the whole epidermo-pilosebaceous unit from the dermis (**Fig. S1**) (**Gilliam *et al.,* 1998**). LCs were then liberated by trypsin digestion and expression of the Ki-67 cell division marker determined by flow cytometry (**Fig. 2B, C**). As expected (**Chorro *et al.,* 2009**), LC proliferation was high at d5 and then declined. Strikingly, we found a reactivation of LC proliferation (22 ± 4.6%) concomitant with early anagen (d27) before the proportion of Ki-67^+^ LCs returned to the low levels of adults (d90). Maximal bromodeoxyuridine (BrdU) incorporation into anagen LCs confirmed the high proliferation rate at this phase of the hair cycle (**Fig. 2D, E**). However, no such anagen-associated increase was observed for dendritic epidermal T cells (DETCs) (**Fig. S2**). Thus, the anagen phase of the natural hair cycle is associated with a high rate of LC proliferation.

**Figure 2:**
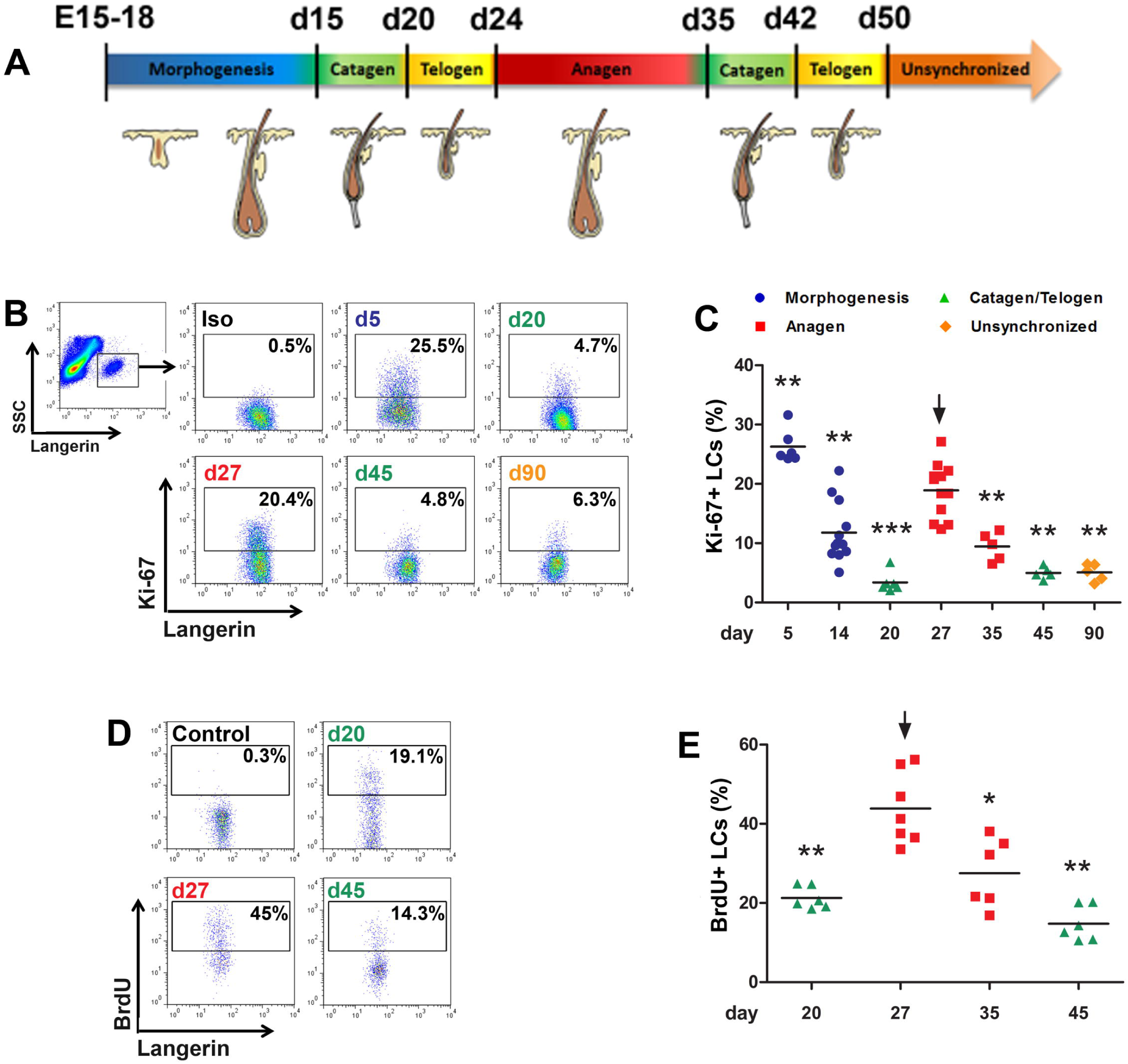
Langerhans cell renewal increases during anagen. **(A)** Schematic representation of the timing of HF morphogenesis and the natural synchronized hair cycle in the back skin of C57BL/6 mice. **(B)** Epidermal cell suspensions from back skin at different hair cycle phases were analyzed by flow cytometry. Langerin+ LCs were gated and their intracellular expression of Ki-67 protein was analyzed. Iso: Ki-67 isotype control antibody labeling performed on d27 mouse skin. **(C)** Percentages of Ki-67^+^ cells among LCs at different time points. Each data point corresponds to one mouse. Data is pooled from 2 independent experiments (d5: n=6, d14: n=12, d20: n=6, d27: n=12, d35: n=5, d45: n=5, d90: n=5). **(D)** Mice were fed or not (control) with BrdU for 4 days. BrdU incorporation into LCs was determined by flow cytometry at the indicated days. **(E)** Percentages of BrdU^+^ cells among LCs at different time points. Each data point corresponds to one mouse. Data is pooled from 2 independent experiments (d20: n=6, d27: n=7, d35: n=6, d45: n=6). Bars correspond to mean values. In **(D)** and **(E),** the arrows indicate the age taken as a reference for statistical analysis (Student’s t-test, *p<0.05, ** p<0.01, *** p<0.001)

### Hair cycle manipulation modifies Langerhans cell proliferation

To further establish a direct relationship between LC proliferation and the growth phase of the hair cycle, we next tested whether physical and genetic manipulations of the cycle would affect LC turnover. Hair shaft removal provoked by depilation triggers synchronized hair growth in adults (**Fig. 3A**) (**Chase, 1954;Paus *et al.,* 1990**). We measured LC proliferation in this model at different time points and observed a peak at early induced anagen (d+7) (**Fig. 3B**).

**Figure 3:**
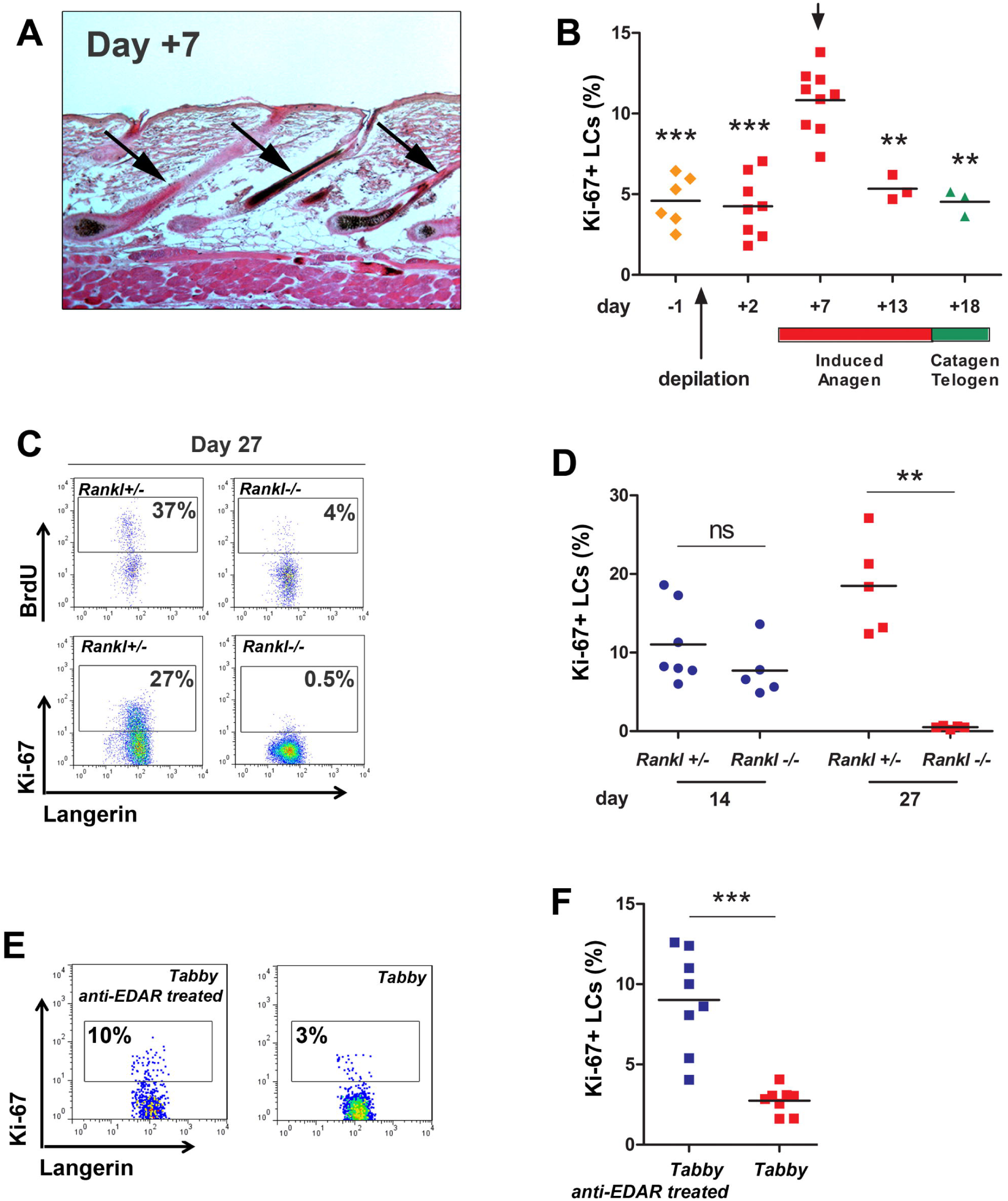
HF cycle manipulation affects the proliferative burst of Langerhans cells. **(A)** Representative image depicting HFs in anagen (arrows) 7 days after depilation. **(B)** Percentages of Ki-67+ cells among LCs in resting phase (d-1), induced anagen (d+2, d+7, d+13) and catagen/telogen (d+18). Data were obtained by flow cytometry on isolated epidermal cells from 2 independent experiments (d-1: n=6, d+2: n=7, d+7: n=9, d+13: n=3, d+18: n=3). In panel B, the arrow indicates the time point taken as a reference for statistical analysis. **(C)** Representative dot plots showing BrdU^+^ and Ki-67^+^ cells among Langerin^+^ LCs in *Rankl*^+/−^ (normal hair cycle) and in *Rankl*^−/−^ skin (no transition into anagen) at the expected time point for anagen (d27). **(D)** Percentages of Ki-67^+^ cells among LCs in *Rankl*^+/−^ and *Rankl*^−/−^ mice during morphogenesis (day 14) (n=7 and n=5, respectively) and at the expected time point for anagen (d27; n=5 and n=5, respectively). Each data point corresponds to one mouse. **(E)** Langerin^+^ LCs from tail skin were analyzed for Ki-67 expression in *Tabby* mice rescued by anti-EDAR agonist antibody injection and untreated, hairless *Tabby* mice. **(F)** Percentage of Ki-67^+^ cells among LCs. Each data point corresponds to one mouse. Data is pooled from 2 independent experiments (n=8 in each group). Bars correspond to mean values. In panels B, D and F, statistical analysis was performed using Student’s t-test (** p<0.01, *** p<0.001).

A well-described side effect of cyclosporin A (CsA), a widely prescribed immunosuppressant drug, is the enhancement of hair growth (**Paus *et al.,* 1998**). We followed LC proliferation in mice shaved then injected subcutaneously with CsA **(**Fig. S3A**).** As expected, the back fur grew back faster in CsA-treated mice **(**Fig. S3B**).** In line with this, the rate of BrdU+ LCs was increased 5-11 days after the CsA treatment **(**Fig. S3C**).**

Next, we investigated mice deficient for TNF-family member RANKL/TNFSF11, which are unable to transit into anagen (**Duheron *et al.,* 2011**). Although LCs retained a normal capacity to proliferate shortly after birth (d14), the anagen proliferation burst observed in control mice (d27) was missing in *Rankl*^−/−^ mice (**Fig. 3C, D**). Finally, we made use of the *Tabby* mice that lack tail HFs. At d35 when tail hair undergoes anagen (**Hodgson *et al.,* 2014**), LC division was measured in *Tabby* mice and compared with *Tabby* mice rescued by embryonic administration of agonist anti-EDAR antibody (**Kowalczyk *et al.,* 2011**). At the expected time for anagen, tail LC proliferation of *Tabby* mice was clearly reduced in comparison to rescued mice (**Fig. 3E, F**). Taken together, these findings demonstrate that anagen is directly associated with a high LC proliferation rate.

### Dividing Langerhans cells are physically associated with the hair follicle

To investigate the spatial relationship between proliferating LCs and cycling HFs, we labelled anagen skin cross-sections for Langerin and Ki-67. Similar to the epidermal sheet overview shown in Figure 1A and in line with previous findings (**Breathnach, 1963;Moresi and Horn, 1997;Christoph *et al.,* 2000**), LCs were found in both the interfollicular areas and the upper portion of the HF (**Fig. 4A**). The number of Ki-67^+^ LCs was determined in the interfollicular section (blue outline) and the HF (yellow outline). We found that the large majority of LCs undergoing cell division resided in or close to the HF (**Fig. 4B**). This finding suggests that the anagen HF conveys LC proliferation signals.

**Figure 4:**
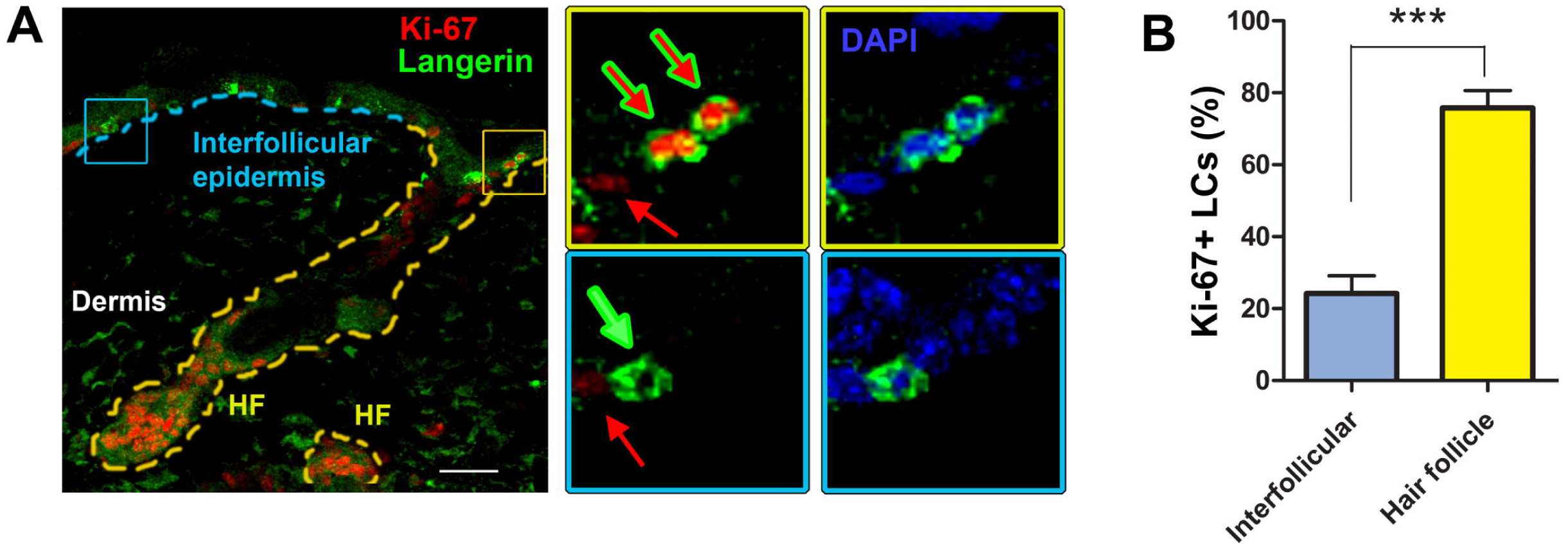
Proliferating Langerhans cells are localized close to the hair follicle. **(A)** Transversal sections of back skin with HFs in anagen were labeled with anti-Langerin (green) and anti-Ki-67 (red) antibodies, counterstained with DAPI (blue) and visualized by confocal microscopy. Two distinct areas were discriminated for analysis: interfollicular epidermis (dashed blue line) and HF (dashed yellow lines). Red arrows indicate Ki-67^+^ epidermal keratinocytes, green arrow indicates LCs and red/green arrows indicate Ki-67^+^ LCs. Pictures are optical slices from Z-stack acquisitions. Scale bar: 50μm. **(B)** Percentages of Ki-67^+^ proliferating LCs within the HF and the interfollicular epidermis. The data is compiled from a total of 50 HFs analyzed from 3 different mice. Statistical analysis was performed using Student’s *t*-test (*** p<0.001).

### Anagen-associated Langerhans cell proliferation relies on CSF-1R signaling

The cytokine IL-34 is expressed by epithelial cells and plays a critical role in the maintenance of the LC network (**Greter *et al.,* 2012;Wang *et al.,* 2012**). Because CSF-1R is the high-affinity receptor for IL-34 (**Lin *et al.,* 2008**), and previous reports have confirmed identical effects of CSF-1R blocking and deficiency in IL-34 (**Greter *et al.,* 2012**), we assessed the functional relevance of CSF-1R signaling on LC renewal. Therefore, we administered antagonistic anti-CSF-1R antibody to mice at the onset of anagen and for 3 days, together with BrdU (**Fig. 5A**). This regimen was sufficient to impair CSF-1R signaling, as shown by a strong reduction of monocytes in peripheral blood **(**Fig. S4A**) (**Greter *et al.,* 2012**)** without disturbing transition into anagen (**Fig. S4B**). Although the treatment did not decrease the overall percentage of LCs in the epidermis (**Fig. S4C**), CSF-1R blocking led to a substantial repression of LC proliferation (**Fig. 5B, C**). In contrast, the cell division of DETCs (**Fig. S4D, E**) and keratinocytes (**Fig. S4D, F**) remained unchanged. This suggests that CSF-1R signaling is implicated in LC proliferation during the hair cycle growth phase.

**Figure 5:**
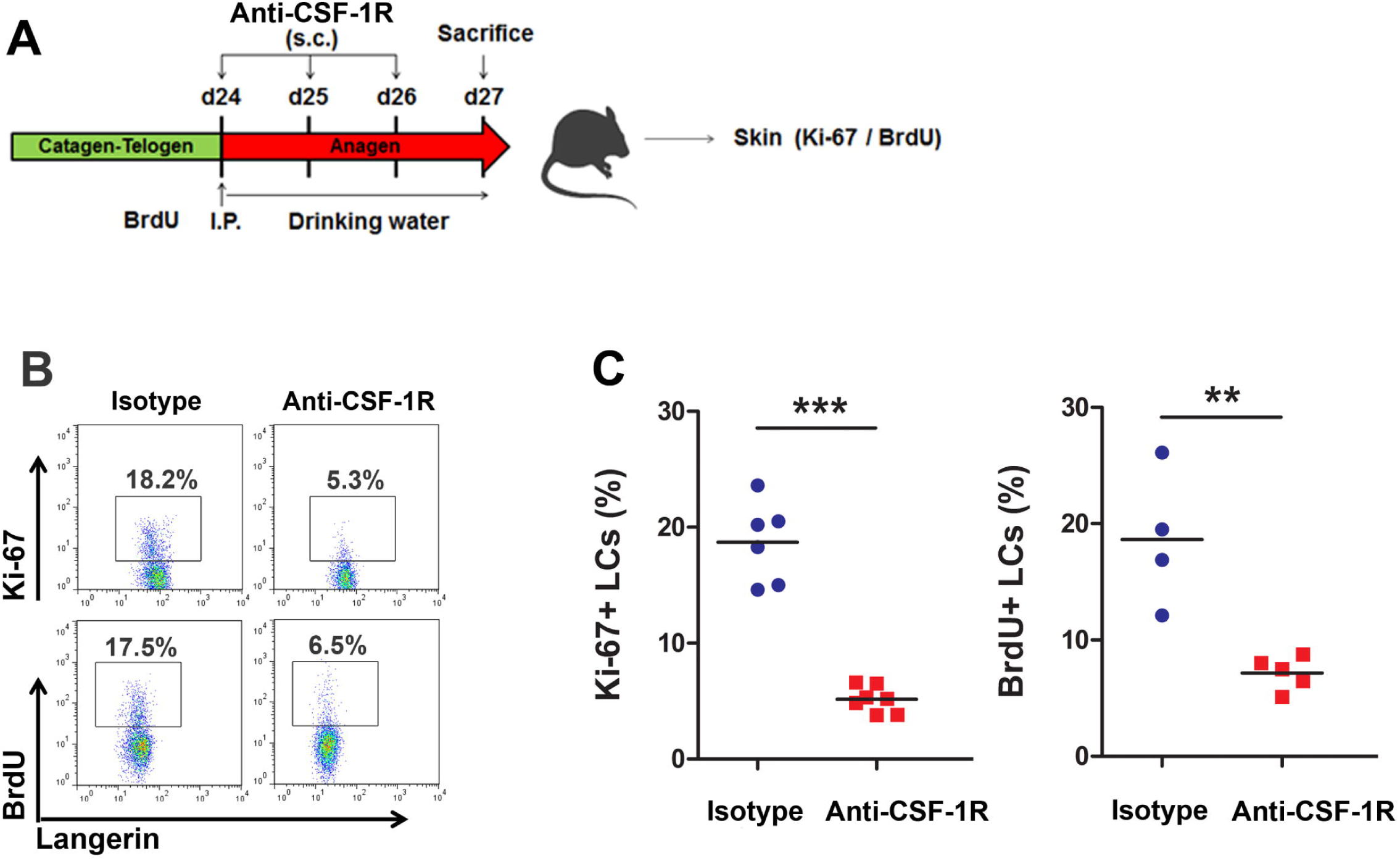
Signaling of IL-34 receptor CSF-1R is required for the anagen-driven Langerhans cell proliferation. **(A)** Schematic representation of the experimental procedure. At the onset of anagen (d24), the mice received a subcutaneous (s.c.) injection of anti-CSF-1R blocking antibody together with BrdU (intraperitoneal [I.P.] and in drinking water). Anti-CSF-1R injections were repeated every other day until sacrifice. **(B)** The percentage of proliferating Langerin^+^ LCs was assessed by Ki-67 expression and BrdU incorporation in mice having received isotype control or anti-CSF-1R blocking antibody. **(C)** Bar graphs show the compiled percentages of Ki-67^+^ or BrdU^+^ cells among LCs for each individual mouse from two independent experiments (BrdU: Iso n=4, CSF-1R n=5; Ki-67: Iso n=6, CSF-1R n=7). Statistical analysis was performed using the Student’s *t*-test (** p<0.01, *** p<0.001).

### IL-34 is expressed by anagen-activated hair follicle stem cells

Since CSF-1R blocking affected LC proliferation in anagen, we addressed the question of whether the HF produces IL-34. Using *Il34*^tm1a^ mice that express the ***LacZ*** gene under control of the *Il34* promoter (**Greter *et al.,* 2012;Wang *et al.,* 2012**), we observed β-galactosidase activity in the interfollicular epidermis and in the upper part of the HF, corresponding to the infundibulum (**Fig. 6A**). No obvious difference could be visualized when comparing HF in anagen vs. telogen.

**Figure 6:**
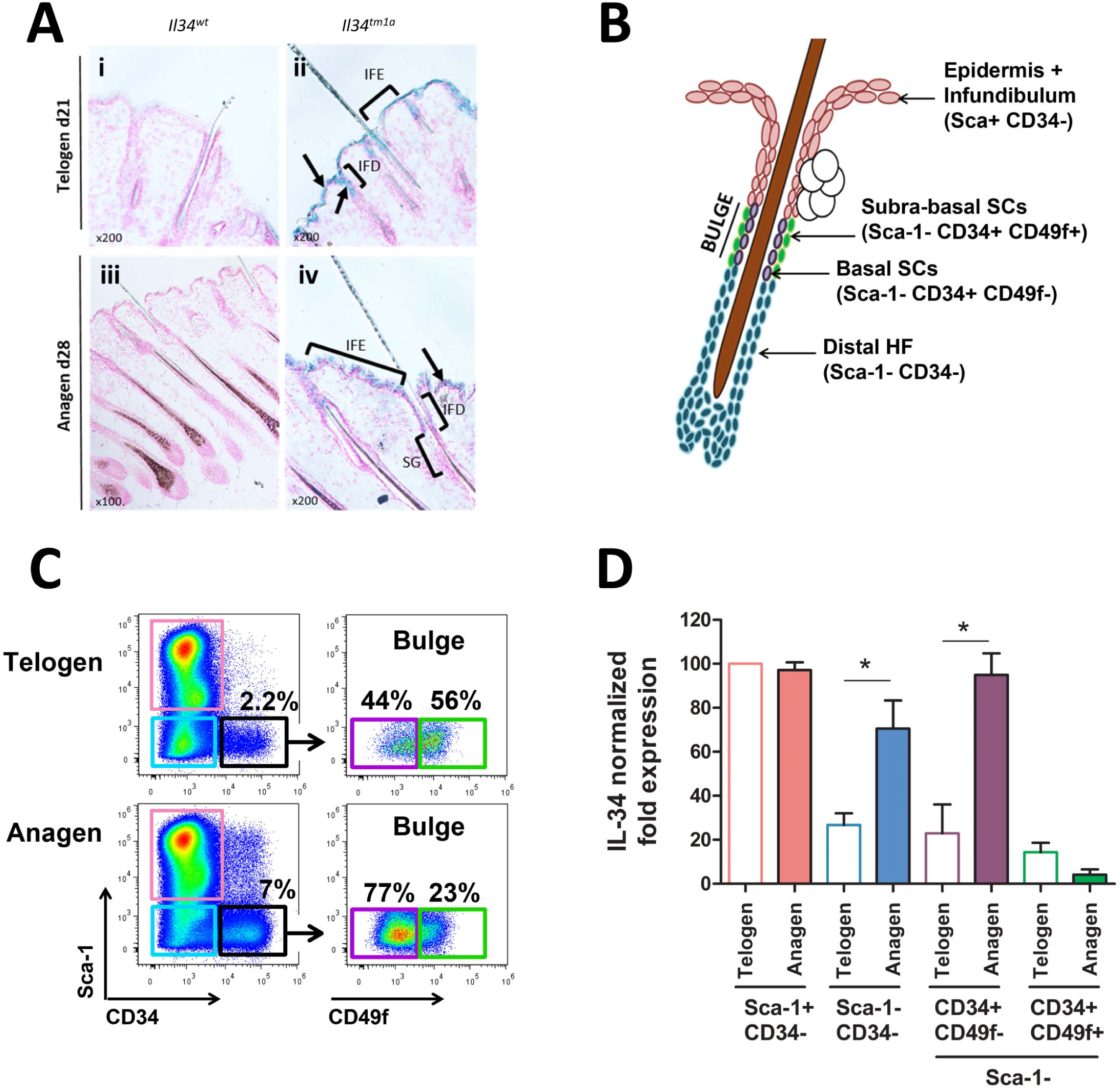
*Il34* expression increases in hair follicle stem cells in anagen. **(A)** Back skin sections (thickness: 16μm) mice were obtained from wild-type (i, iii) or *Il34^tm1a^* (ii, iv) mice during telogen (d21) and anagen (d28) phase. Blue X-gal staining (arrows) depicts areas with LacZ activity. Sections where observed with 200x magnification. IFE: interfollicular epidermis, IFD: Infundibulum, SG: Sebaceous Gland, **(B)** Schematic representation of HF epithelial cell subsets expressing specific marker combinations. The infundibulum and the interfollicular epidermal cells (pink) express Sca-1. Stem cells (SCs) in the bulge are characterized by CD34 expression. As opposed to suprabasal SCs (green), basal SCs (purple) carry CD49⍰/6 integrin. Epithelial cells of the distal HF (blue) lack both Sca-1 and CD34. **(C)** Flow cytometry profile of the CD45^neg^ epithelial cells with color-coded gates corresponding to the subsets defined in panel A. Representative percentages of bulge SCs in telogen and anagen are shown. **(D)** *Il34* mRNA expression by sorted epithelial cell subsets both in telogen or anagen was measured by quantitative RT-PCR and normalized to three housekeeping genes. Bar graphs show the mean − /+ SEM of 3 different mice, expressed as a percentage of telogen epidermal cells. Statistical analysis was performed using the Mann-Whitney test (* p<0.05).

To extend this observation, we sorted the different epithelial cell types of anagen and telogen HFs (**Fig. 6B, C**) (**Jensen *et al.,* 2008;Nagao *et al.,* 2012**) and measured *Il34* transcriptional activity in each subset (**Fig. 5C**). Among non-hematopoietic (CD45^neg^) cells, we first distinguished Sca-1^+^ CD34− interfollicular and infundibulum keratinocytes. For this subset, there was no difference in *Il34* mRNA synthesis between the two phases, similarly to our observations in the IL-34tm1a model (**Fig. 6D**). The Sca-1^−^ CD34^−^ cycling portion of the HF showed an increase in *Il34* mRNA during anagen. An even greater induction of ***Il34*** expression was noted for the CD34^+^ CD49f^−^ stem cells of the bulge area, a cell subset which in addition is amplified during anagen (44 ± 5 % in telogen versus 77 ± 1.5 % in anagen) (**Fig. 6C**). Thus, although this could not be visualized in *Il34*^tm1a^ mice, HF-associated suprabasal stem cells or their progeny might contribute to a localized increase in IL-34 production during hair growth.

## DISCUSSION

The immune sentinel function of LCs implies a fine regulation of their epidermal network. This most likely depends on matching their in situ proliferation with their rate of migration to LNs. Although the cyclic renewal of HF has recognized effects on skin physiology, its impact on LCs had so far not been addressed. In this study, we present evidence that the hair cycle exerts a strong influence on LC self-renewal.

By comparing the rate of cell division between postnatal LC development, the different phases of the hair cycle and adult resting skin, we uncovered a strikingly high level of LC renewal during the first synchronized anagen phase. More precisely, it occurred at early anagen, when the HF is in its maximal activity. Mutants that fail to transit into anagen (*Rankl*^−/−^) or that are devoid of HFs (*Tabby*) lacked this proliferation burst. However, DETCs, a specialized subset of epithelial gamma/delta T cells also residing in the epidermis and capable of self-renewal (**Honjo *et al.,* 1990;Sumaria *et al.,* 2011**), were unresponsive to natural anagen. These findings demonstrate a direct and specific relationship between the hair cycle and LC self-renewal. Moreover, LCs underwent increased cell division in response to synchronized anagen in the adult. depilation is a well-established model to reinitiate hair growth in the adult, when otherwise the hair cycle occurs in a stochastic fashion (**Paus *et al.,* 1998;Plikus *et al.,* 2011**). Although it is difficult to totally rule out some degree of inflammation in this model, it should be noted that anagen-associated LC proliferation occurs one week after depilation, i.e. after the acute inflammatory response. It is therefore probable that the hair cycle also affects LC renewal in the unsynchronized animal, although such measures are offset by the simultaneous occurrence of catagen, telogen and anagen phases in different body areas (**Hodgson *et al.,* 2014**).

CSF-1R is the high affinity receptor for cytokines CSF-1/M-CSF and IL-34 (**Lin *et al.,* 2008**). A number of elements incited us to ask whether the CSF-1R / IL-34 axis plays a functionally important role in anagen LC proliferation. First, the expression of CSF-1R is critical for LC development (**Ginhoux *et al.,* 2006**). Secondly, only the deletion of *Il34*, but not *Csf1*, results in the absence of LCs (**Wang *et al.,* 2012;Greter *et al.,* 2012**). Finally, previous findings on human monocytes support the importance of CSF-1R in cell division (**Clanchy *et al.,* 2006;Lin *et al.,* 2008).** We therefore tested the role of CSF-1R in anagen LC renewal. Detailed analyses of *Il34*^tm1a^ mouse skin showed the presence of *Il34* expression within HFs. The use of antagonistic antibody against CSF-1R (**Greter *et al.,* 2012**) allowed for precisely timed CSF-1R inhibition, thereby restricting the blocking effects to the anagen phase without interfering with development of the LC network (Wang, Greter). This approach revealed the importance of CSF-1R signaling for LC proliferation during anagen. The unresponsiveness of keratinocytes and DETCs to the blocking antibody strongly suggests that CSF-1R blocking has no effect on the other epidermal cell types.

The lack of a role for CSF-1 in LC development and homeostasis (**Ginhoux *et al.,* 2006**), the higher affinity of IL-34 for CSF-1R **(Lin *et al.,* 2008),** and the demonstration that CSF-1 is not expressed by murine epidermal cells (http://biogps.org/#goto=genereport&id=12977) (**Greter *et al.,* 2012**) strongly supports that IL-34 is the CSF-1R ligand responsible for LC proliferation. However, our experiments do not formally exclude a contribution of CSF-1, which has been suggested in the context of cutaneous inflammation (**Wang *et al.,* 2016**). Despite our efforts, we did not succeed in reliably estimating the local concentration of IL-34 protein with commercially available ELISA kits. Although the ***LacZ*** reporter system did not reveal any clear difference of *Il34* promoter activity between anagen and telogen HFs, *Il34* transcription determined by RT-qPCR clearly increased in anagen in the stem cell compartment of the so-called bulge. This region lies just below the constant portion of the HF where reside those LCs that undergo most cell division during anagen. In this context, it can be noted that, in a model of human skin activation by UV, the majority of dividing LCs also localized to the distal part of the HF (**Gilliam *et al.,* 1998**). It was not possible to further dissect the infundibulum from the epidermis because of lack of cell-specific markers.

Previous studies in the context of inflammation suggest that LC renewal and migration to LNs are intrinsically linked (**Katz *et al.,* 1979;Ginhoux *et al.,* 2006**). Our attempts to evaluate LC density at catagen and anagen proved unreliable because the interference of the numerous HF in back skin precluded a clear LC visualization. In addition, the result would have been questionable in the context of a rapidly growing juvenile mouse. Yet, the previous finding that *Rankl*^−/−^ mice display a reduced LC density comes in support of the idea that anagen is required to positively adjust the network density (**Barbaroux *et al.,* 2008**).

Altogether, our study highlights a novel link between the hair cycle and LC homeostasis in steady-state conditions. The demonstration that LC self-renewal is regulated by the hair cycle should incite further investigations into factors released by this ectodermal appendage on the skin immune system.

## MATERIAL AND METHODS

### Mice

Mice were housed in specific-pathogen free conditions facilities at the Institut de Biologie Moléculaire et Cellulaire (Strasbourg, France) and at the Faculté de Biologie et Médecine de Lausanne (Lausanne, Switzerland). Il34tm1a mice were provided by Dr. Frédéric Lezot (Nantes, France). OT-II mice were purchased from Charles River Laboratories (L’Arbresle, France). All experiments were carried out in conformity to the French and Swiss animal bioethics legislation. *Rankl*^−/−^ (**Duheron *et al.,* 2011**) and EDA-deficient *Tabby* (**Gaide and Schneider, 2003**) mice have been previously described. Due to gender-related hair growth kinetics, experimental procedures were performed on male mice only.

### Antibodies and reagents

Anti-Ki67-PerCP-Cy5.5 (clone B56), anti-CD103-PE (M290), anti-CD45-APC (30-F11), anti-CD4-PerCP-Cy5.5 (RM4-5) and anti-CD8α-APC (53-6.7) antibodies were purchased from BD Biosciences (Franklin Lakes, NJ). Anti-TCRγ/δ-ΡΕ (GL3) and anti-Ia/Ie -APC or -PerCP-Cy5.5 (M5/114.15.2) antibodies were purchased from BioLegend (San Diego, CA). Anti-CD11c-PerCP-Cy5.5 (N418), anti-CD45.1-APC or -PerCP-Cy5.5 (A20) and anti-CD4-PE (GK1.5) antibodies were purchased from eBioscience (San Diego, CA). Anti-CD45-PE-Cy7 (I3/2.3) antibody was purchased from Cell LAB (Beckman-Coulter, Brea, CA). Anti-Langerin-AlexaFluor (AF) 488 or −AF647 (929F3.01) antibodies were purchased from Dendritics (Lyon, France). Anti-EpCAM (G8.8) antibody was purchased from Abcam (Cambridge, UK).

### BrdU incorporation

Mice were intra-peritoneally injected with 1mg BrdU (Sigma-Aldrich, St-Louis, MO) in saline per 20-g body weight on the first experimental day. Drinking water was supplemented with 0.8 mg/mL BrdU for 3 or 4 days depending on the experimental settings. Mice were then sacrificed and back skins were collected.

### Skin depilation

Adult mice (>90 days old) were anesthetized by intra-peritoneal injection of 100μg/g body weight of Ketamine mixed with 10μg/g body weight of xylazine. The hair of the back skin was trimmed by an electric razor before removal with cold wax (Klorane).

### CSF-1R blocking in vivo

Mice were anesthetized by isoflurane inhalation and received on the first experimentation day 0.5mg of anti-CSF-1R blocking antibody AFS98 (**Fend *et al.,* 2013;Sudo *et al.,* 1995;Greter *et al.,* 2012**), kindly provided by Dr. Hélène Haegel (Transgene SAS, Strasbourg) or rat IgG2a isotype control (2A3, BioXcell, West Lebanon, NH) by sub-cutaneous injections. Each following day mice were similarly injected with 0.25mg of blocking or isotype control antibodies until sacrifice.

### Reversion of hairless tail phenotype in EDA-deficient *(Tabby)* mice

Briefly, hair follicle deficiency in the tail of *Tabby* mice was reverted by intravenous injection of 100μg anti-EDAR agonist antibody (clone EDAR3) into pregnant mice at E14 of gestation (**Kowalczyk *et al.,* 2011**). Skin cell suspensions. Back or tail skins were taken at precise timings corresponding to HF morphogenesis (d5 and d14), catagen/telogen (d20 and d45), anagen (d27 and d35) or unsynchronized (d90) phases, as previously described **(Lin *et al.,* 2009).** For epidermal cell isolation, the hypodermis was mechanically removed from tail and back skins with a razor. Then skins were incubated, dermal side down, in RPMI medium (Lonza) supplemented with 2% FCS and 1mg/mL dispase II (Roche) overnight at 4°C. Epidermis was removed from the dermis and incubated at 37°C in trypsin solution (TrypLE Select, Life Technologies) for 45 minutes. Cells were liberated by gentle shaking, filtered through 100μm and 40μm cell strainers (BD) to remove epidermal fragments and hair, and washed in saline supplemented in 2% FCS and 0.2mM EDTA (SE buffer).

### Flow cytometry

To label viable cells and block Fc receptors, cell suspensions were pre-incubated for 20 minutes at 4°C with Fixable Viability Dye eFluor780 (eBioscience) and 2μg/mL of anti-CD16/CD32 antibody (clone 2.4G2, BD). Surface staining was done with 1μg/mL of appropriate antibodies diluted in SE buffer for 15 minutes at 4°C and washed twice. For intracellular staining, cells were fixed and permeabilized for 20 minutes at 4°C (Cytofix/Cytoperm buffer, BD), washed and labeled with 1μg/mL appropriate antibodies for 20 minutes at 4°C. BrdU and Ki-67 detection with Flow BrdU or Ki-67 detection kit (BD Pharmingen) were performed according to the manufacturer’s protocol. Flow cytometry acquisitions were performed with a FACS Gallios™ system (Beckman Coulter) and data was analyzed with Flowjo software (TreeStar, Ashland, OR).

### Epidermal sheet preparation and labeling

Tail skin epidermis from C57Bl/6, *Tabby* or control mice were isolated from dermis by dispase II treatment as described above. Epidermal sheets were fixed with cold acetone on ice for 20min. Sheets were then washed in TRIS-buffered saline (TBS) and non-specifics sites were blocked in TBS supplemented with 5% donkey serum for 1 hour at room temperature. Tissues were then incubated with 1μg/mL anti-Langerin-AF488 or 1μg/mL uncoupled anti-Ia antibody (2G9, BD) for 3 hours at room temperature. After washing, sheets were incubated with 0.5μg/mL A555-coupled donkey anti-rat antibody for 1 hour at room temperature. DAPI staining was then realized for 15 minutes at room temperature. Stained tissues were mounted in medium from Dako (Glostrup, Denmark). Images were acquired on a widefield fluorescence microscopy (Axiovert 200M, Zeiss, Iena, Germany) with a lOx or 20x objective (EC Plan-Neofluar, NA: 1.3).

### Histochemistry and image acquisition

Collected samples comprised pieces of juvenile back skin (d27), depilated skin (d+7) and dermis (d27) isolated from epidermis by dispase II treatment, as described above. Samples were fixed in 4% paraformaldehyde for 48 hours at 4°C. Tissues were then included in paraffin. Briefly, the inclusion protocol consisted of a 3 hour incubation in 70% ethanol, 4 hours in 95% ethanol, 16 hours in 100% ethanol, 24 hours in butanol before 48 hours inclusion in paraffin. Tissue was sectioned (8μm) with a microtome (RM2235, Leica, Wetzlar, Germany) and heated overnight at 58°C. Deparaffinization steps comprised incubations in 100% Toluene, 100% ethanol, 95% ethanol, and distilled water.

Sections obtained from depilated skins and skin from mice treated with anti-CSF-1R blocking antibody or control isotype were colored with hematoxylin and eosin. Sections of mouse skin (d27) treated or not with dispase II were colored in Masson’s trichrome.

For immunolabeling, juvenile back skin slices were first deparafinized, followed by a demasking step in boiling 10mM EDTA for 30 minutes. Non-specific sites were blocked in TBS supplemented with 5% donkey serum for 1 hour at room temperature. Tissues were then incubated with 1μg/mL anti-Langerin (M200, Santa Cruz, Dallas, TX) and 20μg/mL anti-Ki67 (TEC-3, Dako) antibodies diluted in TBS with 2% donkey serum for 3 hours at room temperature. After 2 washes in saline, sections were incubated with 0.5μg/mL donkey anti-mouse AF488-coupled secondary antibody (Life Technologies) and 0.5μg/mL donkey anti-rat Cy3-coupled secondary antibody (Jackson ImmunoResearch, West Grove, PA) for 1 hour at room temperature. Tissues were then washed three times and labeled with DAPI for 15 minutes at room temperature before mounting in Dako medium. Z-stack acquisitions were performed with the Axio observer Z1 microscope (Zeiss) equipped with a LSM700 confocal head (Oil-objective 40x, EC Plan-Neofluar, NA: 1.30). Compilations and analysis were realized with Image J software (Macbiophotonics, NIH). For statistics, at least 50 hair follicles were analyzed.

### LacZ stainings

Sections (12μm) of *Il34^tm1a^* and *Il34^wt^* epidermis embedded in OCT were cut using Cryostat Leica CM3050S. Slices were fixed with PFA 1% 5min, rinced with PBS 1x and incubated in Xgal (5-bromo-4-chloro-3-indolyl-beta-D-galactopyranoside) solution overnight at 37°C. Sections were rinced with PBS 1x, left to dry and mounted with EUKITT® medium.

### Cyclosporin A treatment

C57BL/6 mice were shaved (d-4), then injected subcutaneously with 50μL saline or cyclosporin A (Sandimmun; Novartis) diluted at 50mg/kg into each flank. Injections were repeated at d-2 and d0. At d5, d11 and d17, mice were treated with BrdU as described above. Three days later, epidermal suspensions were generated from back skin and LCs were tested for BrdU incorporation by flow cytometry.

### FACS sorting, RNA extraction and quantitative RT-PCR

Isolated epidermal cells from telogen (d20) or anagen (d27) skin were sorted on a FACSAria II (BD) on the basis of their Sca-1, CD34 and CD49f profile expression. Sorted cells (>95% purity upon post-sort verification) were lysed in RLT buffer (Qiagen, Venlo, The Netherlands). RNA was extracted with RNeasy Micro kits (Qiagen), and cDNA was synthesized with oligo(dT)15 primers and the Thermo Scientific Maxima First Strand cDNA Synthesis Kit. Quantitative PCR was performed using Thermo Scientific Luminaris Color HiGreen qPCR Master Mix and ran on a Stratagene MX4000 thermal cycler. Gene-specific primers are listed in **Table S1**. CT values of target genes were normalized to GAPDH, HPRT and β-actin. The expression factor was calculated by using the Relative Expression Software Toll (REST, http://mmm.gene-quantification.de/rest.html).

## ACKNOWLEDGEMENTS

The authors would like to thank Astrid Hoste for assistance with immunohistochemistry, Monique Duval and Delphine Lamon for mouse care, Dr. Jean-Daniel Fauny for his help with confocal microscopy analyses and Claudine Ebel (Institut de Génétique et de Biologie Moléculaire et Cellulaire, lllkirch, France) for cell sorting. We also thank Dr. Hélène Haegel (Transgene, Strasbourg, France) who provided anti-CSF-1R blocking antibody. We are indebted to Pr. Nikolaus Romani for critical reading of the manuscript as well as Pr. Ralf Paus for discussion.

This work is supported by the Centre National pour la Recherche Scientifique and the Agence Nationale pour la Recherche (Program “Investissements d’Avenir”, ANR-10-LABX-0034 MEDALIS; ANR-ll-EQPX-022).

B. Voisin is funded by the French Ministry of Research and Higher Education and by the Fondation pour la Recherche Médicale (FDT20130928345). V. Flacher and C.G. Mueller are supported by the CNRS and European Union grants (Marie-Curie Career Integration Grant “Dermacro PCIG12-GA-2012-334011” to V. Flacher and Internal Training Network “STROMA ITN-289720” to C.G. Mueller).

## CONTRIBUTIONS

B. Voisin, D. Chartoire, C. Devouassoux and C. Kowalczyk-Quintas performed experiments. F. Clauss and P. Schneider provided *Tabby* mice. F. Lezot provided IL-34tmla mice. B. Voisin, V. Flacher and C.G. Mueller wrote the manuscript. The authors have no conflicting financial interests.

## ONLINE SUPPLEMENTAL MATERIAL

**Figure S1:**
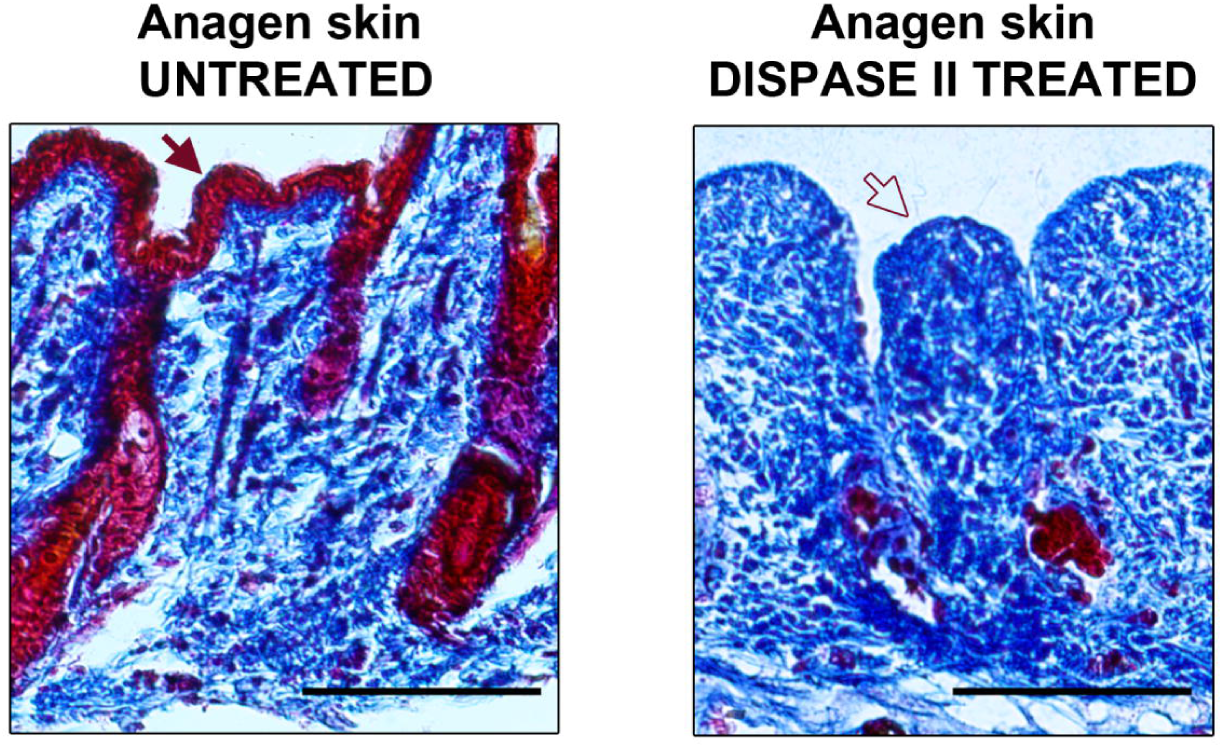
Dispase II efficiently isolates the epidermis together with hair follicles. Anagen back skin was treated (right panel) or not (left panel) with dispase II by floating skin onto culture medium containing the enzyme overnight at 4°C. Separation of epidermis from dermis was performed on treated skin, which was then fixed and included in paraffin. Transversal sections were colored in Masson’s trichrome. Filled red arrow indicates the epidermis in untreated skin while empty red arrow shows its absence after dispase II treatment. Scale bar: 100μm.

**Figure S2:**
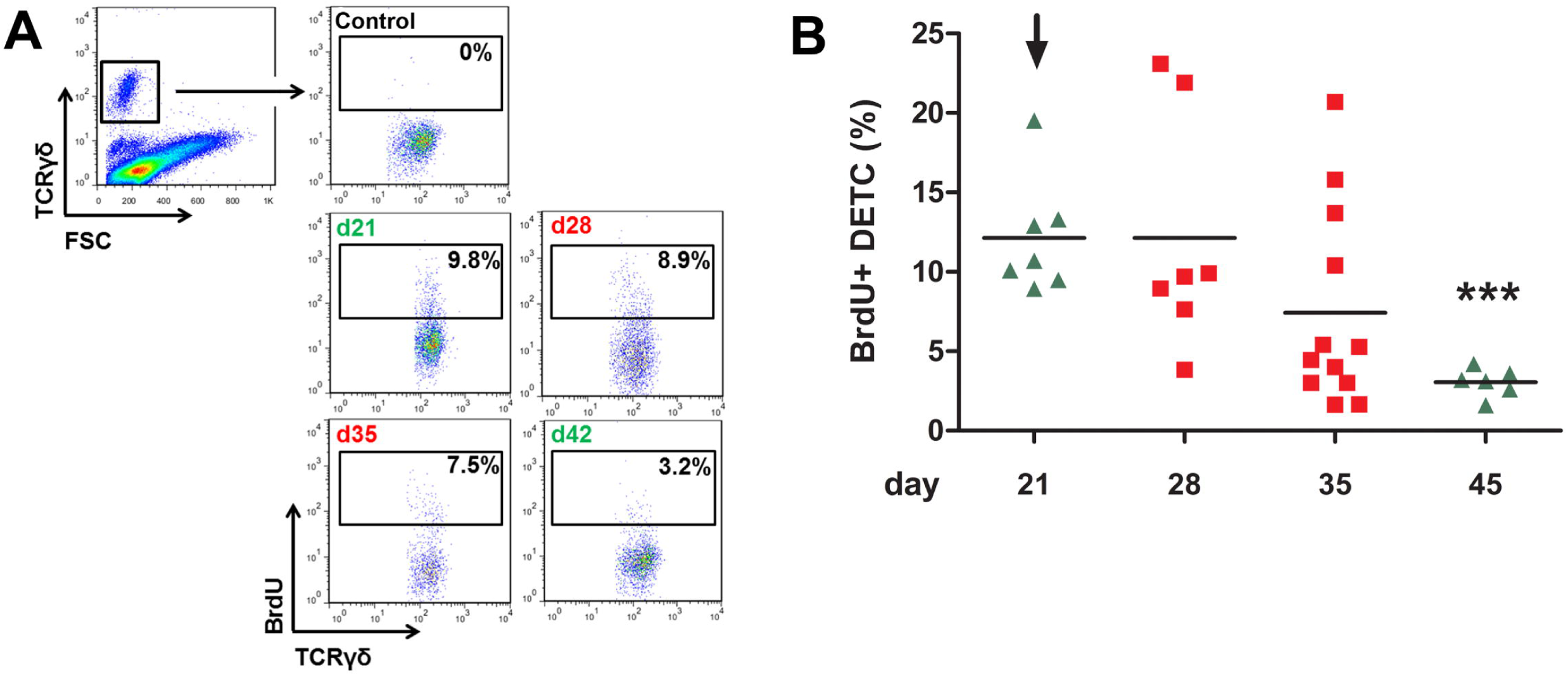
Proliferation of dendritic epidermal T cells does not depend on the natural hair cycle. **(A)** Mice were fed or not (control) with BrdU for 4 days and its incorporation into DETCs (TCR⍰δ^+^) was measured by flow cytometry at the indicated days. **(B)** Percentages of BrdU^+^ cells among DETCs. Data is pooled from 2 distinct experiments (d21: n=7, d28: n=7, d35: n=12, d45: n=6). Bars represent mean values. The arrow indicates the age taken as a reference for statistical analysis (Student’s t-test, *** p<0.001).

**Figure S3:**
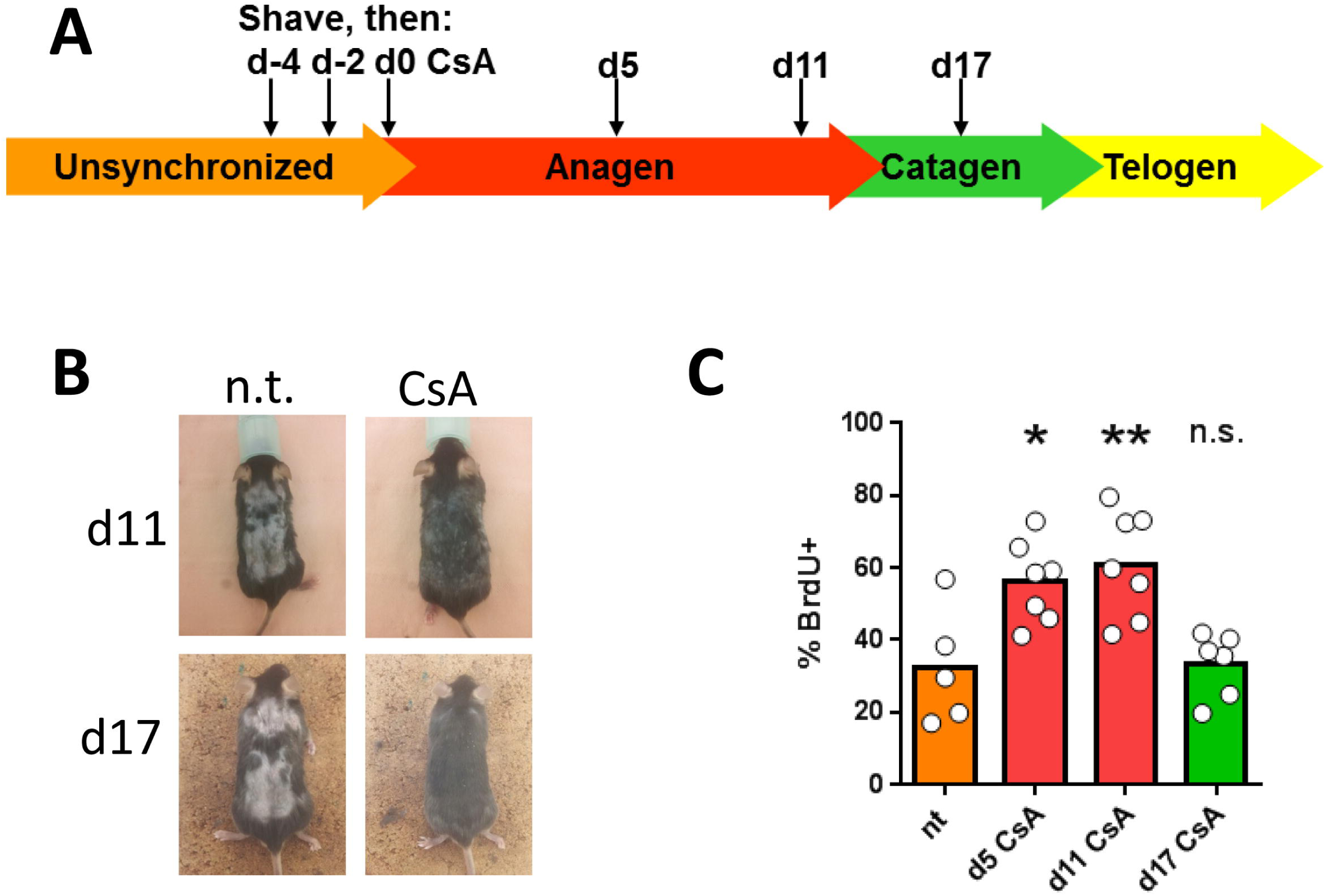
Cyclosporin A promotes hair growth and LC proliferation. Mice were shaved at d-4 and treated with cyclosporin A (CsA) at d-4, d-2, d-0. **(A)** Representative pictures of the back skin at d11 and d17 for untreated and CsA-treated mice. **(B)** BrdU incorporation in epidermal LCs 5, 11 or 17 days after CsA treatment.

**Figure S4:**
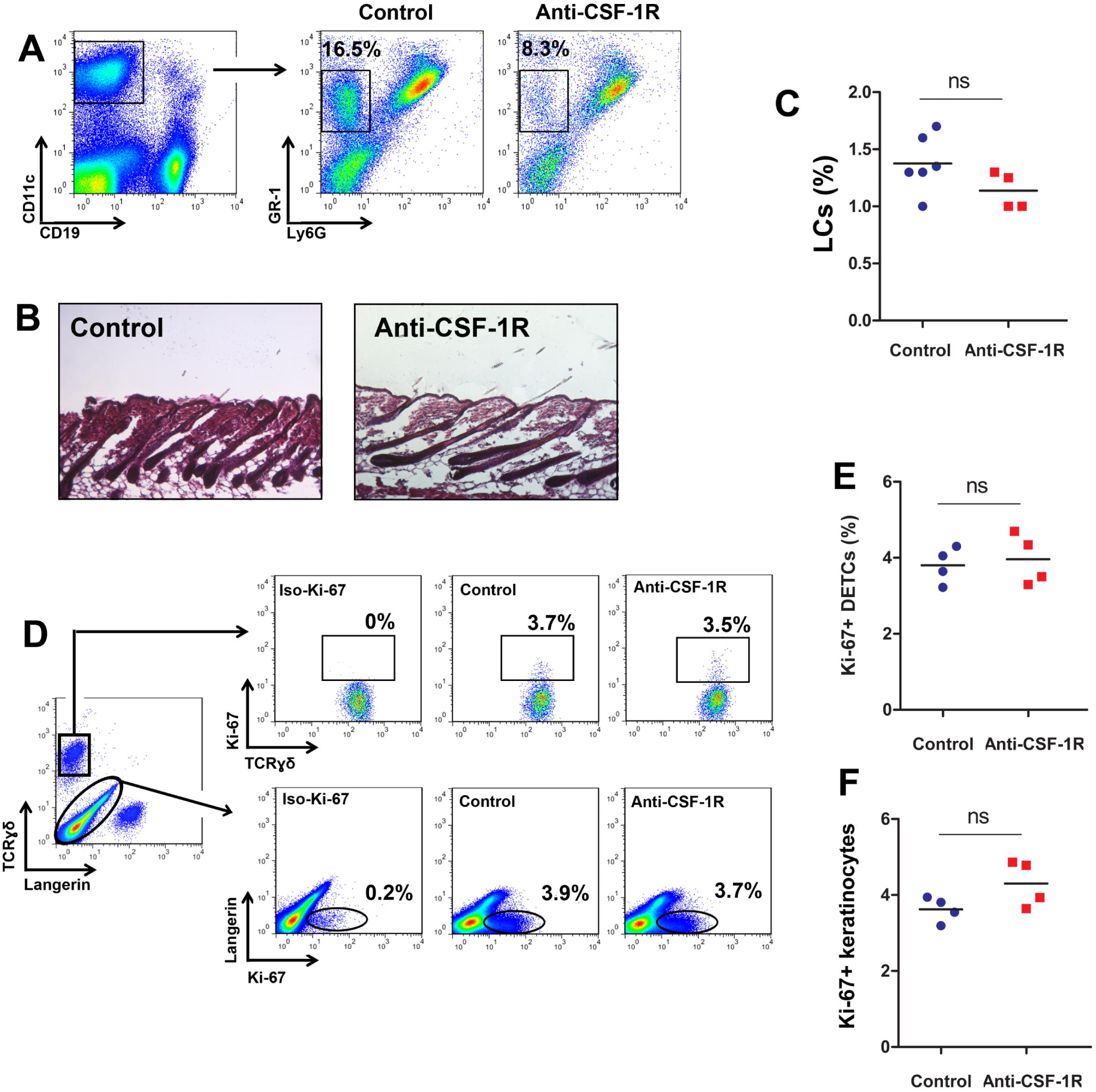
Transient blocking of CSF-1R does not affect the hair cycle nor the proliferation of keratinocytes or dendritic epidermal T cells. Mice received subcutaneous injections of anti-CSF-1R or isotype control (control) from day 24 to day 27. **(A)** Blood was collected at day 27 to assess the efficiency of CSF-1R blocking by the loss of monocytes (CD11b^+^ GR-1^+^ Ly6G-) in injected mice as compared to control. **(B)** Entry into anagen in treated or control mice skin at day 27 was determined by hematoxilin eosin colorization of back skin slices. **(C)** Percentage of LCs (Langerin+) was determined in epidermal suspension of treated (n=4) or control mice (n=6) skin. Data were obtained from 2 distinct experiments for each group. **(D)** Proliferation of DETC (TCR⍰δ+ cells) and keratinocytes (Langerin-TCR⍰δ-) was analyzed by the expression of Ki-67 in skin epidermal suspension of anti-CSF-1R or isotype treated mice. Iso-Ki-67: Isotype control of Ki-67 antibody. **(E)** and **(F)** Graphs show the compilation of flow cytometry results obtained from panel D (n=4 for each group). Data were obtained from 2 distinct experiments for each group. Statistical analysis was performed using the Student’s *t*-test (ns: non significant).

**Supplementary Table 1.**
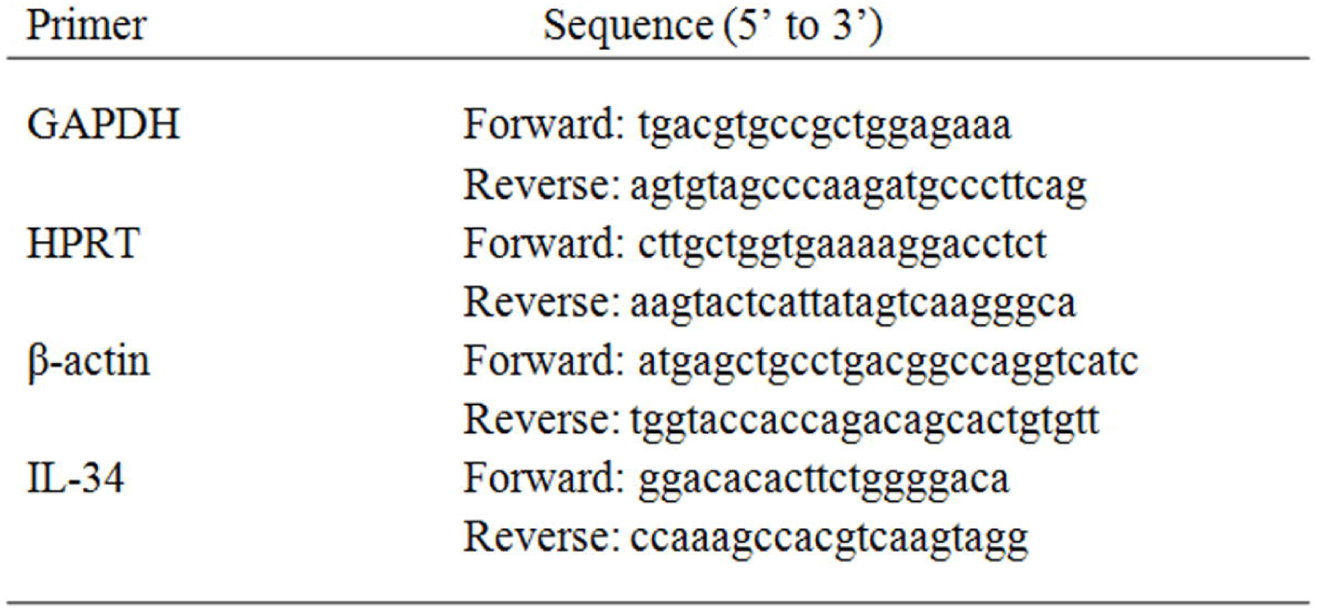

